# An introduced crop plant is driving diversification of the virulent bacterial pathogen *Erwinia tracheiphila*

**DOI:** 10.1101/345009

**Authors:** Lori R. Shapiro, Joseph N. Paulson, Brian J. Arnold, Erin D. Scully, Olga Zhaxybayeva, Naomi Pierce, J. Rocha, Vanja Klepac-Ceraj, Kristina Holton, Roberto Kolter

## Abstract

*Erwinia tracheiphila* is the causal agent of bacterial wilt of cucurbits, an economically important phytopathogen affecting few cultivated Cucurbitaceae host plant species in temperate Eastern North America. However, essentially nothing is known about *E. tracheiphila* population structure or genetic diversity. To address this shortcoming, a representative collection of 88 *E. tracheiphila* isolates was gathered from throughout its geographic range, and their genomes were sequenced. Phylogenomic analysis revealed three genetic clusters with distinct *hrp*T3SS virulence gene repertoires, host plant association patterns, and geographic distributions. The low genetic variation within each cluster suggests a recent population bottleneck followed by population expansion. We showed that in the field and greenhouse, cucumber (*Cucumis sativus*), which was introduced to North America by early Spanish conquistadors, is the most susceptible host plant species, and the only species susceptible to isolates from all three lineages. The establishment of large agricultural populations of highly susceptible *C. sativus* in temperate Eastern North America may have facilitated the original emergence of *E. tracheiphila* into cucurbit agro-ecosystems, and this introduced plant species may now be acting as a highly susceptible reservoir host. Our findings have broad implications for agricultural sustainability by drawing attention to how worldwide crop plant movement, agricultural intensification and locally unique environments may affect the emergence, evolution, and epidemic persistence of virulent microbial pathogens.

**Importance:** *Erwinia tracheiphila* is a virulent phytopathogen that infects two genera of cucurbit crop plants, *Cucurbita* spp. (pumpkin and squash) and *Cucumis* spp. (muskmelon and cucumber). One of the unusual ecological traits of this pathogen is that it is limited to temperate Eastern North America. Here, we complete the first large-scale sequencing of an *E. tracheiphila* isolate collection. From phylogenomic, comparative genomic, and empirical analyses, we find that introduced *Cucumis* spp. crop plants are driving the diversification of *E. tracheiphila* into multiple, closely related lineages. Together, the results from this study show that locally unique biotic (plant population) and abiotic (climate) conditions can drive the evolutionary trajectories of locally endemic pathogens in unexpected ways.

## Introduction

Complex interactions between human behavior, demographic change, the local environment and microbial evolution underlie the emergence and transmission of the pathogenic microorganisms that have shaped human history. Many pathogens first emerged into human populations during the Neolithic Revolution, when the widespread adoption of agricultural technologies led small, isolated hunter-gather groups to settle into larger, denser civilizations. These agriculture-driven demographic changes facilitated the emergence and evolution of some virulent microbial pathogens that specialized on humans as hosts (1-3). These newly emerged, human-specialized pathogens remained geographically restricted until global trade and human migration inadvertently introduced these pathogens to novel geographic areas (4, 5). Despite modern advances in medicine and public health, complex local ecological conditions – such as exponential human population growth, rapid urbanization, human-livestock and human-wild animal contact, and microbial evolution – are continuing to drive local emergence of novel human pathogens (6, 7). Devising strategies to predict pathogen emergence, and to control newly emerged pathogens remains one of the most pressing public health concerns, and, deservedly, is attracting intense international research efforts (8).

Less recognized is that similarly complex anthropogenic and ecological factors are likely driving the emergence of microbial pathogens into cultivated crop plant populations. Humans are continually creating new ecological niches by transforming complex ecological habitats into simplified agro-ecosystems (1, 9-11). Since the Neolithic Revolution, and accelerating with global trade, the geographic range of many crop plant species has expanded from the limited geographic region of origin (where the wild progenitors evolved with the endemic biotic communities for millions of years) to worldwide cultivation (12, 13). This creates landscapes of crop plants with distinct biogeographic histories suddenly being cultivated in close proximity to each other, and at times to wild, undomesticated progenitors. These mosaic landscapes facilitate continual encounters between locally endemic insects and microbes, with high density populations of genetically similar native and introduced crop plants (14-18). This increases the probability that a novel virulent pathogen will be generated through mobile DNA invasion, and subsequently encounter a large, genetically homogeneous host population into which it can emerge and then rapidly spread.

*Erwinia tracheiphila* Smith (Enterobacteriaceae), the etiological agent of bacterial wilt of cucurbits, is one plant pathogen with genomic changes consistent with a recent emergence into a novel host plant population (19, 20). *E. tracheiphila* is a highly virulent phytopathogen only known to affect two genera of Cucurbitaceae crop plants – *Cucumis* spp. (cucumber and muskmelon) and *Cucurbita* spp. (pumpkin, squash, and yellow-flowered gourds). *E. tracheiphila* induces characteristic wilt symptoms by blocking xylem sap flow (Figure 1), causing infected plants to die within 2-3 weeks after the first symptoms appear. Curiously, losses due to *E. tracheiphila* are only reported from a very limited geographic range in temperate Midwestern and Northeastern North America (20-30). This conspicuously contrasts with the worldwide distribution of susceptible cucurbit host plant populations (31-34). *Cucurbita* spp. are native to the New World, and undomesticated *Cucurbita* populations naturally occur from subtropical South America through the Southeastern United States (35-37). Wild *Cucumis* ssp. are native to the Eurasian, Australian and African tropics and subtropics (33). *Cucumis* spp. did not occur in Eastern North America until Spanish colonists introduced cultivated varieties in the early 1500s (33, 38). *E. tracheiphila* causes the most severe losses in introduced *Cucumis* spp. crop plants, and less severe losses in native *Cucurbita* spp. crops (25, 29, 39, 40). *E. tracheiphila* is obligately insect-transmitted by two species of highly specialized leaf beetle species that are endemic to Eastern North America: the striped (*Acalymma vittatum*) and spotted (*Diabrotica undecimpunctata*) cucumber beetles (Coleoptera: Chrysomelidae: Luperini: Galerucinae: Diabroticina). *E. tracheiphila* transmission can occur when frass from infective beetles contacts recent leaf wounds or floral nectaries (25, 41-45). Direct losses from leaf beetle herbivory and *E. tracheiphila* infection, and indirect costs to control populations of the beetle vector amount to many millions of dollars annually (29).

**Figure 1.**
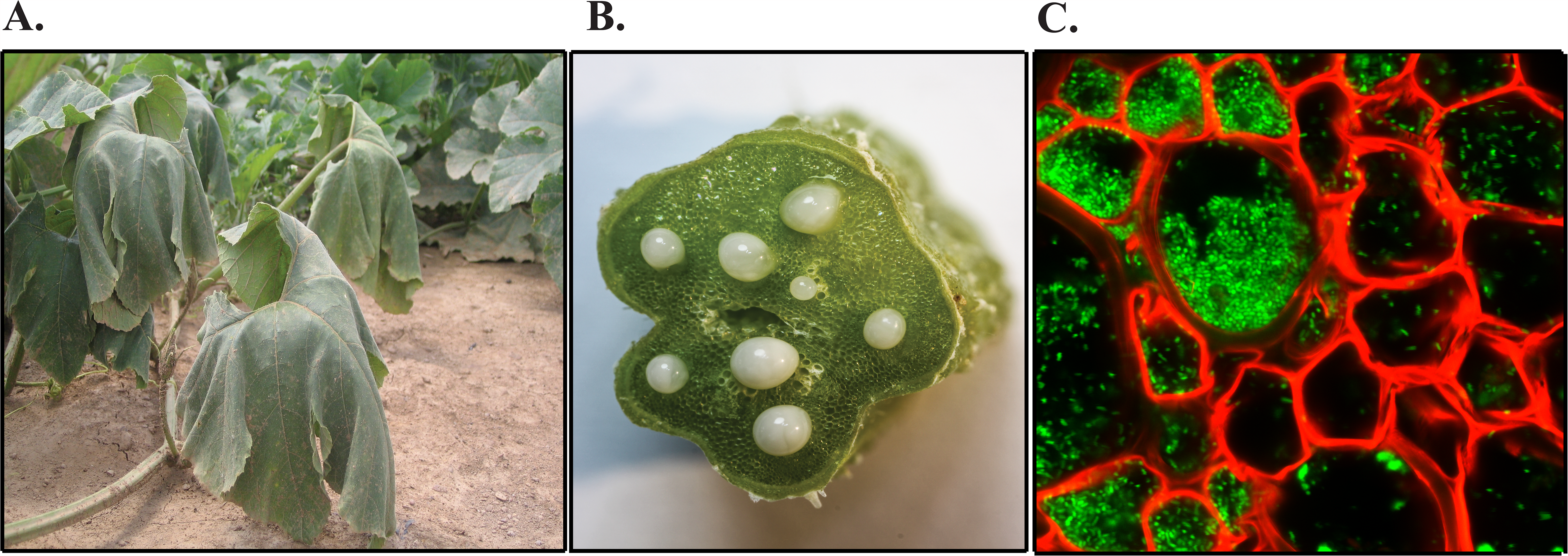
*Erwinia tracheiphila* infection at the macroscopic and microscopic levels. **A.** A vine of a field-infected *Cucurbita pepo* plant shows characteristic systemic wilting symptoms. **B.** *E. tracheiphila* can be seen oozing from multiple blocked xylem vessels in a cross-section of a symptomatic cucumber stem. **C.** *In planta* confocal microscopy image of *E. tracheiphila* (green) blocking the xylem (red) of a wilting squash plant.

Despite its economic burden, nothing is known about the population structure of *E. tracheiphila*, the genetic basis of virulence against the two cucurbit genera that *E. tracheiphila* infects, or why *E. tracheiphila* only occurs in such a restricted geographic range. To address this knowledge gap, we collected and sequenced an 88 isolate collection sampled from all susceptible host plant species across the entire geographic range where *E. tracheiphila* is known to occur. Via analysis of the genomes of these isolates, we evaluated *E. tracheiphila* genetic diversity in relation to its plant host range and geographic distribution. We find that these isolates group into three distinct clusters that differ in host plant associations, geographic ranges and horizontally acquired virulence gene repertoires. Low genetic heterogeneity and an excess of rare alleles within each lineage are consistent with a recent bottleneck and expansion into a susceptible host population. We then tested for interactions between the abiotic environment (temperature) and host plant species on *E. tracheiphila* virulence in controlled inoculation experiments. We find that *E. tracheiphila* is more virulent at temperate compared to subtropical summer temperatures. Further, we find that cucumber – a crop plant recently introduced into Eastern North America – is the most susceptible to *E. tracheiphila* overall, and the only plant species susceptible to infection by isolates from all three lineages. From this, we infer that both genetic factors (i.e., horizontal acquisition of virulence genes) and ecological factors (i.e., foreign crop plant introductions and low genetic diversity in agricultural populations) may have driven the recent emergence and epidemic persistence of *E. tracheiphila* into cucurbit agricultural populations in temperate Eastern North America.

## Results

### *Erwinia tracheiphila* is comprised of three phylogenetic lineages that have different plant host and geographic ranges

Of 88 isolates, 68 were recovered from the introduced *Cucumis* spp. crop plants (cucumber and muskmelon) and only 20 were recovered from native *Cucurbita* spp. crop plants (squash and pumpkin) (Table 1). A phylogenetic network analysis, which can account for and visualize phylogenetic conflict due to recombination and gene flow (46, 47), revealed that *E. tracheiphila* is comprised of three distinct, co-existing phylogenetic clusters, designated as *Et-C1*, *Et-C2* and *Et-melo* (Figure 2a, Supplemental Figure 1). Faint reticulations along the long branches connecting *Et-C1* and *Et-melo* suggests some limited gene flow between these two groups. *Et-C2* is on a non-reticulating long branch and shows no evidence of gene flow with either *Et-C1* or *Et-melo* (Figure 2a). We refer to these three distinct groups as phylogenetic ‘clusters’ instead of ‘pathovars’, as ‘pathovar’ assignments are often inconsistent with phylogenetic group (48-51).

**Figure 2.**
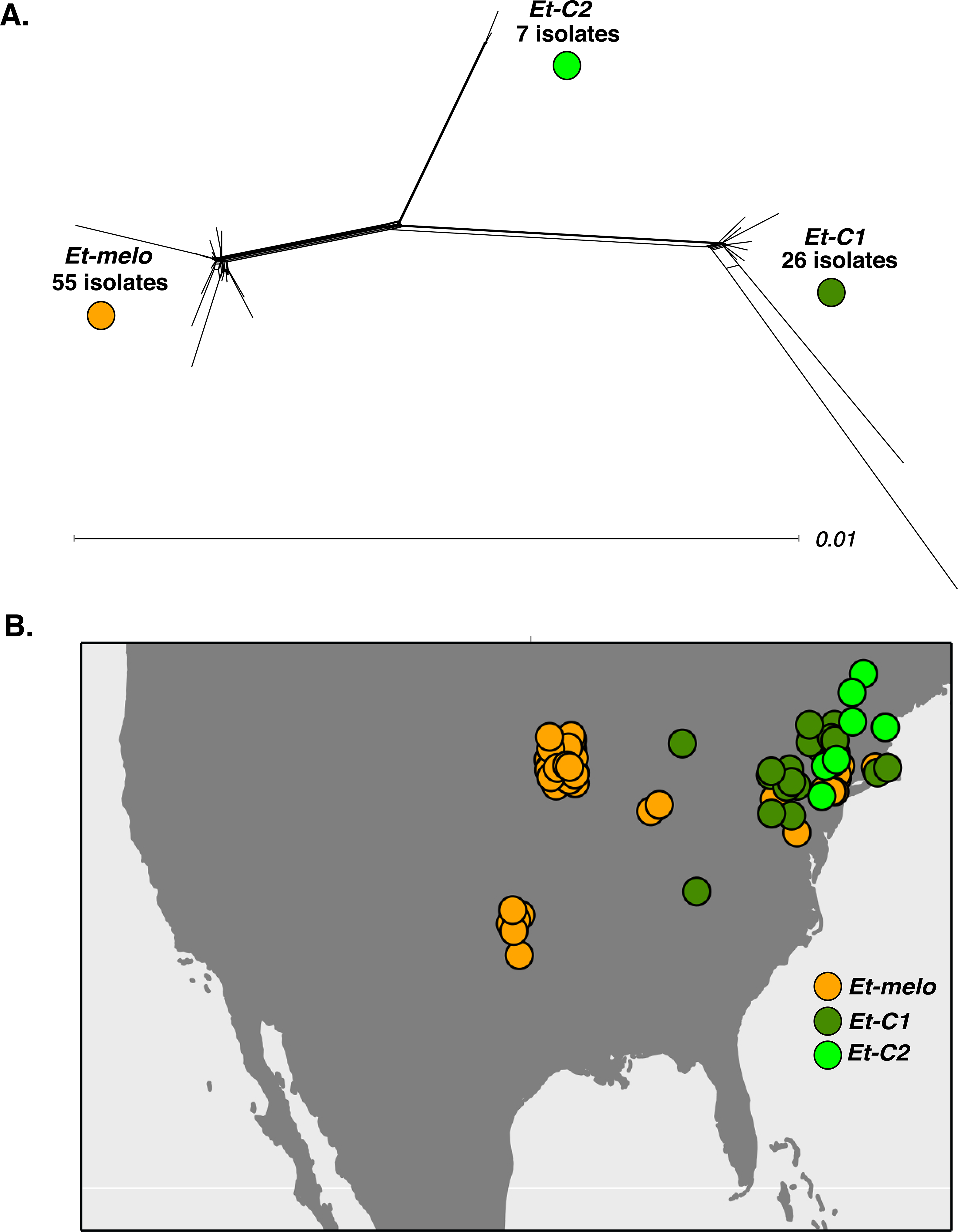
Three genetically distinct lineages of *Erwinia tracheiphila* and their geographic distribution. A. The phylogenetic network of 88 *Erwinia tracheiphila* isolates. The network was reconstructed from concatenated alignments of the 1567 core gene families identified with OrthoMCL in all 88 *E. tracheiphila* genomes. Three distinct clusters separated by long branches are named as *Et-melo*, *Et-C1* and *Et-C2* based on the host plant they were found to infect (Table 1): Isolates from clusters *Et-C1* and *Et-C2* were found only on squash and cucumber plants, while strains from cluster *Et-melo* were only found on muskmelon and cucumber. Host plant, year of isolation, location, and assembly metadata for each isolate is listed in the Supplemental Table 1. Scale bar, substitutions per site. Supplementary Figure 1 shows individual isolate IDs for each isolate in the network. **B. Geographic distribution of the three clusters.** Each of the 88 isolates is plotted as a single circle on the map according to its collection site, and colored according to the genetic cluster to which it belongs (see panel A). The isolate-specific locations and year of collection are listed in Supplemental Table 1.

The three clusters are present at different frequencies, over different geographic ranges, and have distinct host plant association patterns (Figure 2b, Table 1, Supplemental Table 1). The most frequently recovered *E. tracheiphila* (55 isolates) belong to the *Et-melo* cluster and were collected exclusively from the cucumber and muskmelon. *Et-melo* also has the largest geographic distribution, encompassing the known range of *E. tracheiphila* throughout the Midwestern and Northeastern United States (Figure 2b). The 26 *Et-C1* isolates were recovered from both introduced cucumber and native squash plants collected in the Mid-Atlantic and Northeast (Table 1). Of the 7 *Et-C2* isolates, six were recovered from squash and one from cucumber (Table 1), and all *Et-C2* isolates were found in the Northeastern United States (Figure 2b). Isolates from all three clusters were found in field infected cucumber plants, while muskmelon was only infected by the *Et-melo* isolates, and squash were infected only by the *Et-C1* and *Et-C2* isolates (Table 1). All three lineages are geographically restricted to temperate Eastern North America (Figure 2b). This is further north than where wild, undomesticated *Cucurbita* spp. naturally occur in the American tropics and subtropics (31, 35, 52).

### All three *Erwinia tracheiphila* lineages have low genetic diversity

To investigate the recent population history of *E. tracheiphila*, genetic diversity was measured with the Watterson estimator (*θ_W_*) and Tajima’s D. These were calculated separately within each phylogenetic cluster and within each collection period (2008-2010 and 2015). The core genes shared by all isolates within each lineage were assigned as putatively functional *(‘Intact’*), or mobile DNA/putatively pseudogenized *(‘Pseudogenized + Repetitive’)* using published, manually-curated gene annotations from the BuffGH reference genome (formerly PSU-1) (20, 30). There is low within-cluster nucleotide diversity (*θ_W_*) in all three lineages (Table 2) despite clear between-cluster genetic divergence (Figure 2a), which is consistent with small effective population sizes. *Et-C2*, which was only observed in the 2015 collection, has the fewest segregating sites, is represented by the fewest isolates in the smallest geographic range, and has isolates with the shortest branch lengths (Figure 2a), which together suggest *Et-C2* may be the most recently emerged lineage (Tables 1 and 2, Figure 2b). For both *Et-C1* and *Et-melo, θ_W_* increased over the 7-year period, although diversity increased 7-fold faster in *Et-C1* than *Et-melo*. The low overall heterogeneity within each *E. tracheiphila* cluster is compatible with recent emergence from a small founder population, and recent divergence into distinct genetic clusters.

In addition to the density of polymorphic sites (*θ_W_*), the allele frequencies at these sites also contains information about recent population history. Tajima’s D, which measures the degree to which the allele frequency spectrum is compatible with that of a neutral population of constant size, is negative for all three clusters (Table 2). This reflects an excess of rare variants and suggests these three lineages are experiencing an ongoing population expansion after a bottleneck. The excess of rare alleles is consistent with the hypothesis that these three relatively monomorphic lineages are rapidly spreading within genetically homogenous host plant populations that are susceptible to infection by pathogen variants with the same virulence alleles. All three lineages show evidence of limited within-lineage recombination, although the large number of repetitive regions likely make recombination estimates inexact (Table 2). While estimated rates of homologous recombination are relatively low for all three clusters, this process may be contributing to lack of within-cluster phylogenetic structure (Figure 2a).

### Estimation of the *Erwinia tracheiphila* core genome, pangenome and functional repertoire

The entire *E. tracheiphila* pangenome of the 88 strains sequenced here, encompassing all core, accessory, and unique genes, is 10,598 gene families (Figure 3a). The pangenomes of geographically widespread microbes with diverse environmental reservoirs such as *Prochloroccocus* or *Escherichia coli* have almost an order of magnitude more gene clusters (53, 54). The relatively small *E. tracheiphila* pangenome size of ~10,600 genes is compatible with the hypotheses that *E. tracheiphila* is a host-restricted pathogen that recently emerged from a population bottleneck and/or is predominantly circulating in low-diversity agricultural host plant populations (20).

**Figure 3.**
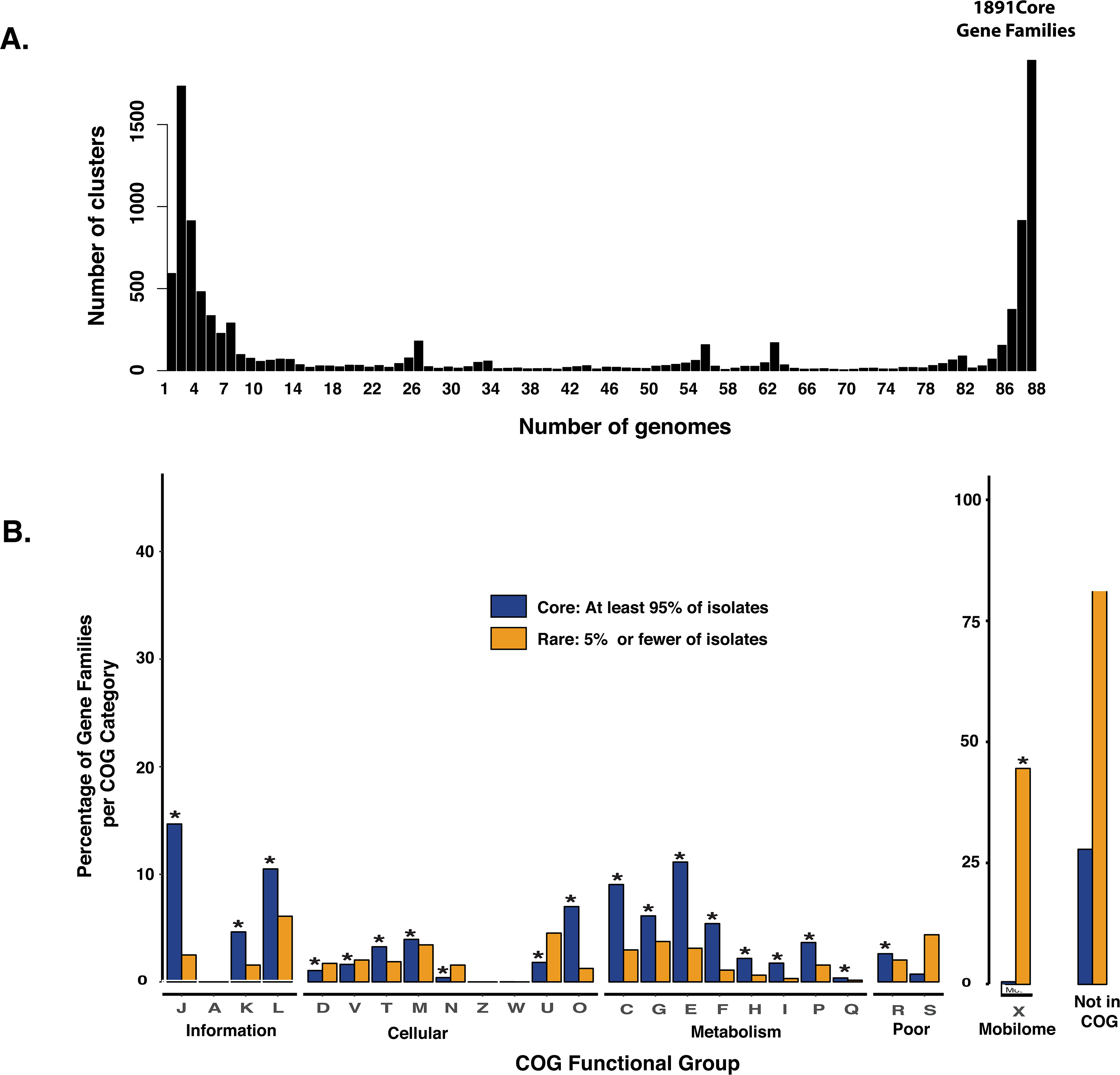
The *Erwinia tracheiphila* pan-genome and its functional annotations. **A**. **Distribution of detected gene families among core and rare pangenome.** The number of gene families (y axis) is plotted as a histogram, with counts of the number of sequenced *E. tracheiphila* genomes that contain them (x axis). Of 10,598 gene families in the species, micropan identifies 1,891 gene families present in all 88 genomes. There are 4,032 gene families present in at least 95% (84 out of 88 genomes) and are referred to as ‘core’ genes. 4,890 gene families are found at intermediate frequency (between 6-84 genomes), and 3,720 gene families are present in less than 5% (6 genomes) and are referred to as ‘rare’ genes **B. Distribution of predicted functions in the core and rare gene families of *E. tracheiphila***. The ‘core’ and ‘rare’ gene families are grouped into COG categories (x axis), which are annotated by their one-letter abbreviations (see Supplemental Table 1 for notations). Y axis shows the percent of the gene families within each COG category. The bar to the far right shows the overall percentage of the core and rare gene families that were not represented in COG. ‘Mobile’ (X) and the number of genes not assigned to a COG are shown with a 100% y-axis, while the other categories are shown with a y-axis scaled to 40%. Asterisks designate the functional categories that are significantly over-represented when compared to the distribution of all genes in that category (Fisher’s exact test, *P* < 0.05; Supplemental Table 2). The percentage of ‘rare’ and ‘core’ genes not in COG (far right) are shown for scale, but are were not included in the statistical tests.

Of the 4,032 gene families present in at least 95% of sequenced genomes, 2,907 (72%) can be assigned to a functional category of the Clusters of Orthologous Groups (COG) database (55). These ‘core’ gene families are enriched in almost all COG categories associated with cellular processes and metabolism (Figure 3b, Supplemental Table 1). This finding is consistent with these gene families being essential for survival, and therefore ubiquitous in all isolates in the population. Only 699 out of 3,720 (18.8%) genes found in less than 5% of *E. tracheiphila* sequenced genomes are assignable to a COG functional category. This set of gene families that are ‘rare’ in the population are only enriched in ‘Mobilome’ (X), suggesting that most rare genes are accessory genes or mobile DNA and are not involved in cellular, metabolic, or information processes (Figure 3b, Supplemental Table 2).

### *Erwinia tracheiphila* clusters vary by *hrp*T3SS effector content

Many Gram-negative bacterial phytopathogens use a <u>hy</u>persensitive <u>r</u>esponse and pathogenicity Type III Secretion System (*hrp*T3SS) to translocate effector proteins directly into the host plant cell. In the plant cell cytoplasm, T3SS effectors may reveal the presence of a pathogen and initiate a cascade of anti-pathogen defenses, often mediated through responses induced by salicyclic acid (56). Alternatively, effectors may promote pathogen virulence by suppressing induced plant defense responses. *E. tracheiphila* contains an *hrp*T3SS locus, and *E. tracheiphila* suppresses salicylic acid production in a wild gourd host (*Cucurbita pepo* ssp. *texana*) (20, 44), suggesting that *E. tracheiphila* may use effectors for suppressing plant induced defenses during disease development.

We found that the 88 *E. tracheiphila* isolates collectively encode at least 23 *hrp*T3SS effector genes (Figure 4, Supplemental Figure 2). Because differences in T3SS effector repertoire can drive host plant specificity, we also examined the distribution of effector genes between the three *E. tracheiphila* clusters. Cluster *Et-melo* has one unique effector gene, Eop3, which is homologous to the Eop3 gene in *Erwinia amylovora* (57), the uncharacterized *P. syringae* pv. *actinidiae* effector HopBN1 (16) and the *P. syringae* effector HopX1 (58). Two other effector genes, NleD and AvrRpm1, are unique to the *Et-C1* cluster. In the BuffGH reference genome, NleD is present in six copies, including in an intact phage region (20, 30). The *E. tracheiphila* NleD genes have 99% amino acid identity to an NleD gene in an active phage region in the emerging mouse pathogen *Citrobacter rodentium* (59) (Supplemental Figure 3). The functional significance for *E. tracheiphila* having six NleD copies if there is functional significance is not yet known. There are no effector genes that are unique only to the *Et-C2* cluster, but a gene for effector HopAM1 is present in *Et-C2* and *Et-C1* isolates, and a gene for HopAF1 is present in *Et-C2* and *Et-melo* isolates (Figure 4). In *P. syringae*, HopAM1 manipulates abscisic acid-mediated responses and water availability via stomatal closure (60), but how it affects the virulence phenotype for *E. tracheiphila* is unknown. In *P. syringae*, HopAF1 inhibits PAMP-mediated increases in ethylene production, and homologs are widely distributed in many bacterial phytopathogens (61). All five cluster-specific effectors (HopAM1, NleD, AvrRpm1, Eop3, and HopAF1) are physically located far from the *hrp*T3SS locus, and their evolutionary histories are all consistent with horizontal acquisition (Supplemental Figure 3). Phytopathogen effectors are often determinants of host range, and the horizontal acquisition of these five effectors may underlie the split of *E. tracheiphila* into phylogenetic clusters with distinct virulence phenotypes and host plant association patterns.

**Figure 4.**
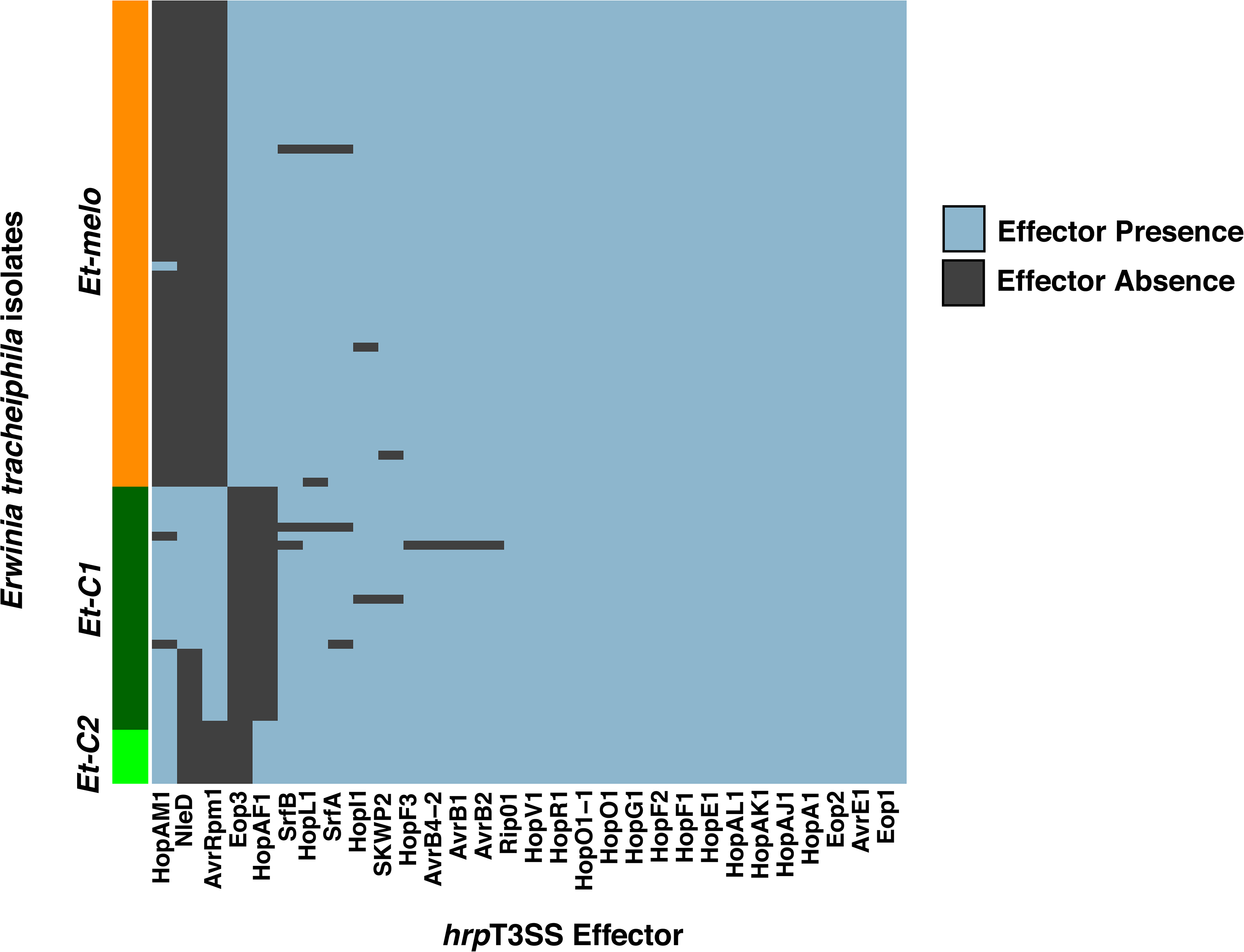
Distribution of *hrp*T3SS effector genes across the genomes of 88 *Erwinia tracheiphila* isolates. Each individual sequenced isolate is represented by a row, and the rows are grouped by phylogenetic cluster (y axis). The 23 effector genes found in the *Erwinia tracheiphila* pangenome are arranged on the x axis. Each cell in the matrix is color-coded by the presence (blue) or absence (dark grey) of a *hrp*T3SS effector gene in a genome of an individual isolate. The effector presence/absence matrix with isolate names is included in Supplemental Figure 2. Phylogenetic trees for the five cluster-specific effectors (HopAM1, NleD, AvrRpm1, Eop3, and HopAF1) are shown in Supplemental Figure 3.

### Cucumber is the only host plant susceptible to all *Erwinia tracheiphila* lineages

Controlled cross-inoculation experiments were used to test whether the patterns of lineage-specific host plant associations observed in the field were due to random sampling patterns, or were reflective of genetic differences. In the greenhouse, three isolates from *Et-melo*, three isolates from *Et-C1*, and one isolate from *Et-C2* were all cross-inoculated into two-week-old seedlings of squash, cucumber, and muskmelon. Isolates from *Et-melo* killed all experimental cucumber and muskmelon plants (Figure 5). In squash, *Et-melo* isolates induced localized wilt symptoms, but all squash plants inoculated with *Et-melo* recovered (Figure 5). Isolates from *Et-C1* and *Et-C2* were highly virulent against cucumber, killing 98% of experimental cucumber plants, but less virulent against squash and muskmelon (Figure 5, Table 3). The attenuation of *Et-C1* and *Et-C2* virulence on muskmelon compared to *Et-melo* in the greenhouse is likely ecologically important, as none of these strains have yet been isolated from field infected muskmelon (Table 1). Squash showed variable susceptibility to isolates from *Et-C1* and *Et-C2*, which is consistent with previous reports that this genus is moderately resistant to *E. tracheiphila* (Figure 5, Table 3) (25). In summary, cucumber is the most susceptible of the three host plant species, and is the only host plant susceptible to infection by isolates from all three *E. tracheiphila* clusters in both the field (Table 1, Figure 2a) and greenhouse (Figure 5).

**Figure 5.**
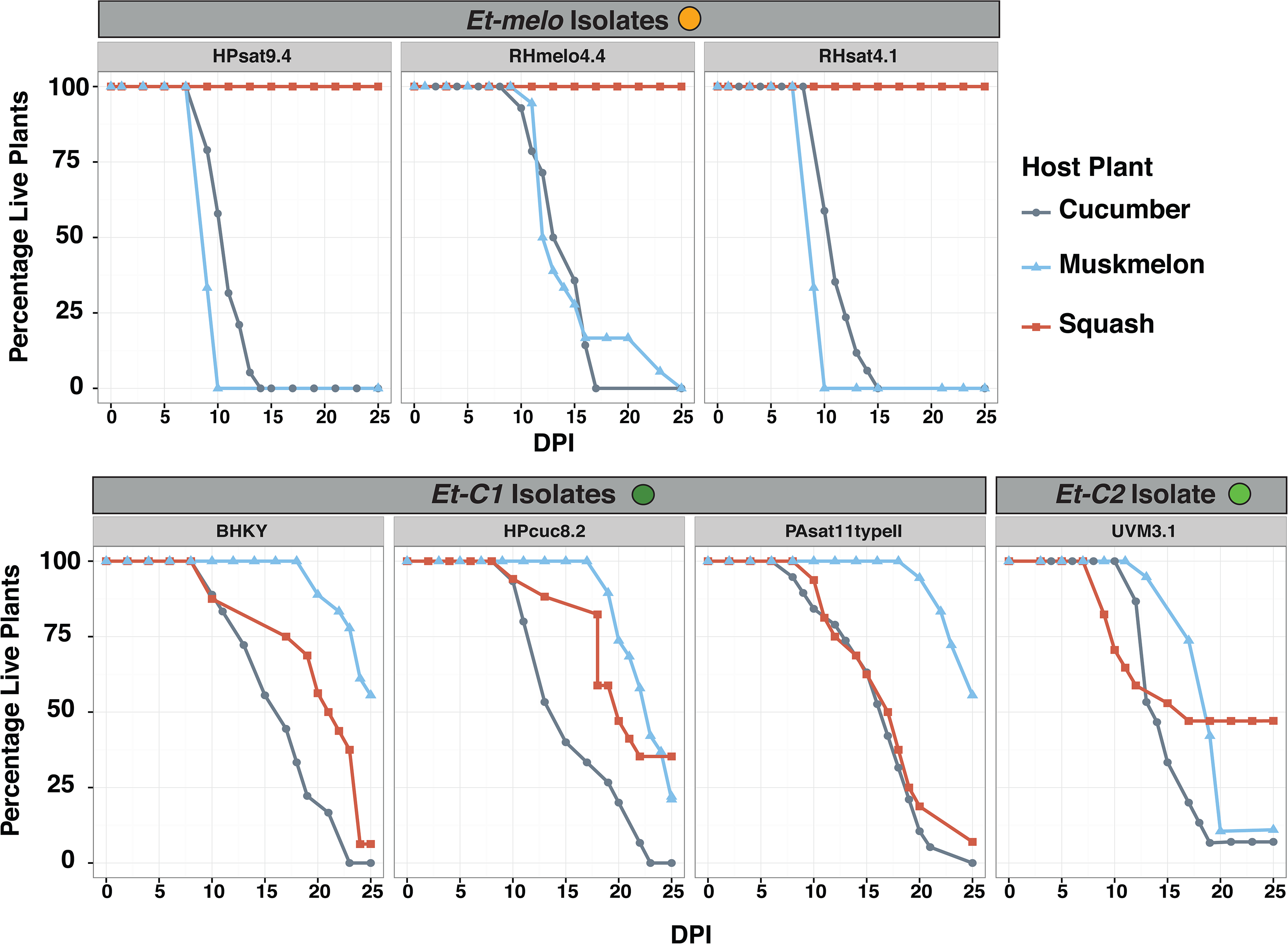
*In planta* virulence of isolates from different clusters in muskmelon, cucumber, and squash. Individual panels show the change in the percentage of live plants (y-axis) over 25 Days Post Inoculation (DPI; x-axis) in controlled greenhouse cross-inoculation experiments. The name of each tested isolate is shown inside a light gray bar, and the isolates are grouped according to the phylogenetic cluster to which they belong (see Figure 2a).

### Sub-tropical temperatures inhibit *Erwinia tracheiphila in vitro* growth and *in vivo* virulence

We tested the effects of temperature on *in vitro* growth and *in vivo* virulence to determine whether the temperatures in temperate Eastern North America, the only region in the world where *E. tracheiphila* is known (see ‘Confirmation of restricted *Erwinia tracheiphila* geographic range’ section in Methods), is more favorable for *E. tracheiphila* growth compared to subtropical temperatures. For isolates from all three clusters, we find that the final OD_600_ concentration after 40 hours of *in vitro* growth is suppressed at warmer 33°C and 37°C incubation temperatures, compared to incubation at cooler incubations at 28°C or 30°C (*P* ≤0.001) (Figure 6, Table 4).

**Figure 6.**
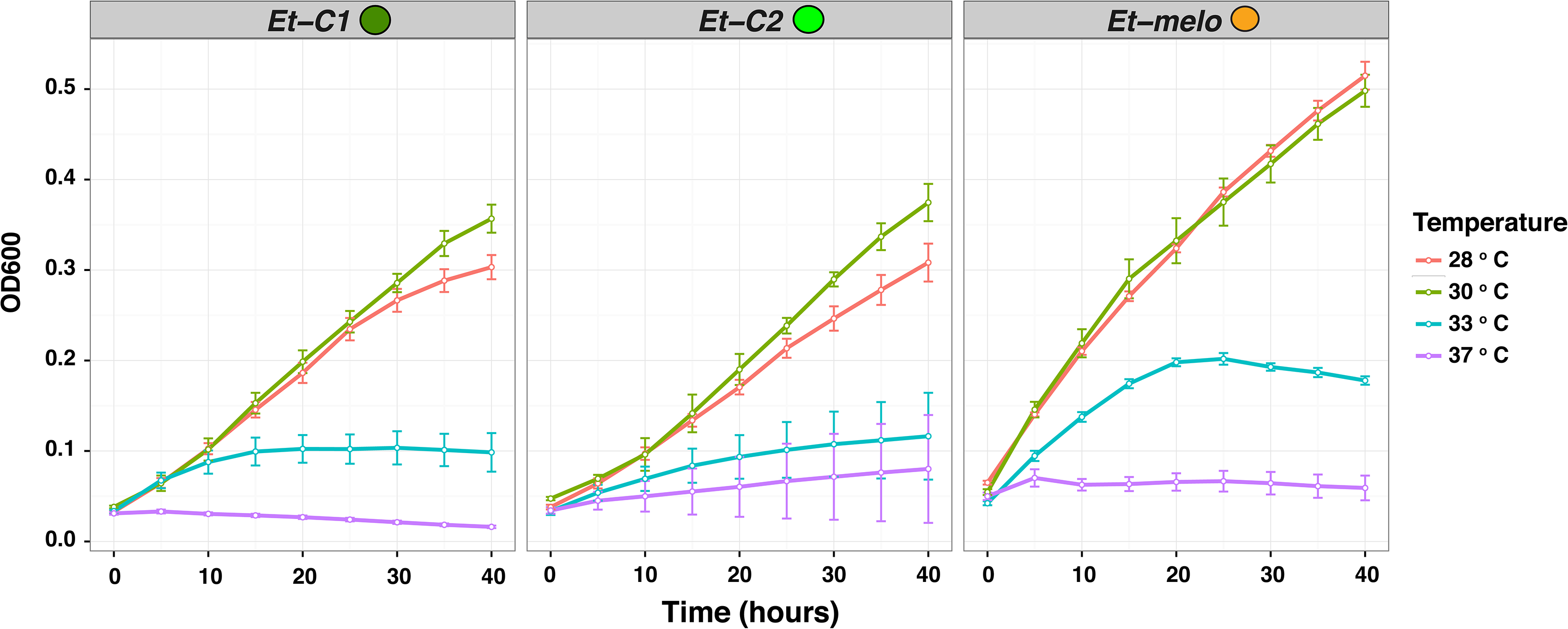
Effects of temperature on *in vitro* growth of *Erwinia tracheiphila.* Individual panels show *in vitro* growth for 7 isolates from *Et-C1*, 2 isolates from *Et-C2*, and 4 isolates from *Et-melo* grown at four different temperatures. Bacterial growth was assessed via optical density (OD_600_; the y-axis), which was measured hourly over 40 hours, and displayed in intervals of 10 hours on the x-axis. Each individual curve shows the average values of all tested isolates in a corresponding cluster. Error bars, standard error of the mean.

To test the effects of temperature on *in vivo* virulence, we isolated an *E. tracheiphila* strain from a field-infected cucumber, and a second strain from a field-infected squash. Each isolate was then inoculated into the host species in which it was found. Half of the plants were incubated at average July temperatures measured in Massachusetts (27°C day/18°C night) to represent the temperate Northeastern United States. This is the region where *E. tracheiphila* is an annual epidemic, all three *E. tracheiphila* lineages were sampled and cultivated cucurbits are only present due to human agriculture. The other half of the inoculated plants were incubated at average July temperatures measured in Texas (33 °C day/23°C night) to represent the subtropical Southwestern United States, where the wild squash progenitor (*Cucurbita pepo* ssp. *texana*) is native but *E. tracheiphila* has never been reported (35). At ‘Southwestern US’ temperatures, only three inoculated squash plants developed localized symptoms in the inoculated leaf, and these three plants recovered. At cooler ‘Northeastern US’ temperatures, half of the squash plants developed localized wilt symptoms, but only six of these plants developed systemic disease and died within the 25-day experiment (Figure 7 and Table 5). In contrast to squash, at ‘Southwestern US’ temperatures, 34 out of 36 cucumber plants died by 21 days post infection (DPI). At cooler ‘Northeastern US’ temperatures, all 36 cucumber plants died by 19 DPI. Moreover, plant death at temperate Northeastern US temperatures occurred significantly faster in cucumber (mean of 13.8 days) compared to squash (mean of 18.3 days). In summary, cucumber is significantly more susceptible than squash at both tested temperatures. Cooler temperatures (normal in the Northern introduced range) are required for *E. tracheiphila* virulence in squash and also increase virulence of *E. tracheiphila* against cucumber.

**Figure 7.**
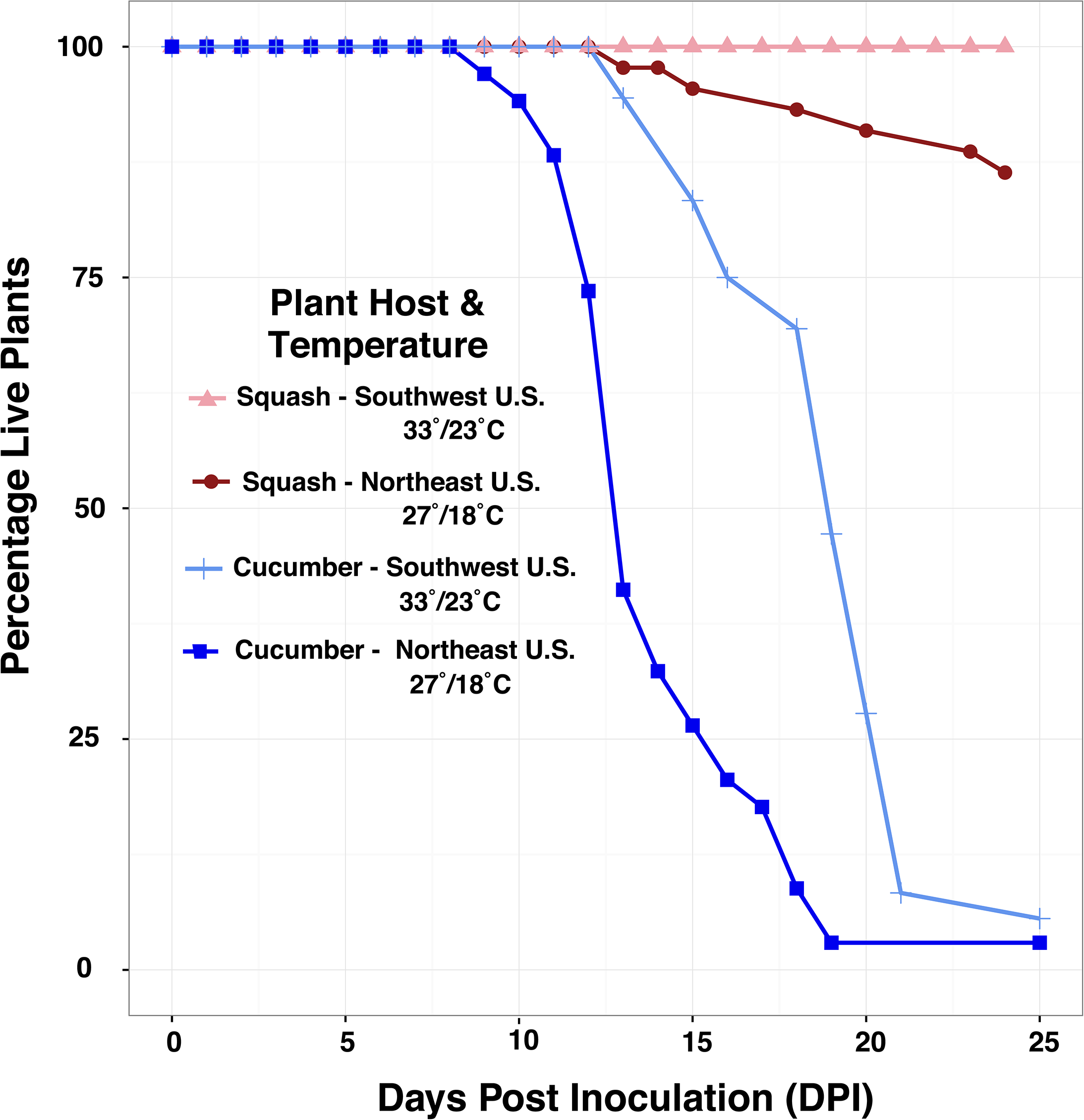
Effects of temperature on *in planta* virulence. Change in percentage of live squash and cucumber seedlings (y-axis) was tracked at two different temperatures over time (x-axis). The time is shown in days since inoculation of plants with *Erwinia tracheiphila* (Day 0). Incubation was at either temperate (dark red for squash and dark blue for cucumber) or subtropical (light red for squash and light blue for cucumber) temperatures.

## Discussion

In our comprehensive study of *Erwinia tracheiphila* genomic diversity, host plant association patterns and demographic history we found that *E. tracheiphila* is comprised of three distinct, homogeneous phylogenetic lineages that have an excess of rare genetic variants. From this, we infer that these three clusters were recently founded by small populations, and are currently experiencing rapid population expansions to fill new agro-ecological niches (3, 70, 82, 86). These inferences about *E. tracheiphila* demographic history correlate with recent anthropogenic changes to cucurbit agro-ecosystems in Eastern North America. The recent introduction of all cucurbit crop plants into temperate Eastern North America, one of the world’s most agriculturally intensive regions, likely created a novel ecological niche (33, 64, 65). Cucumber, a recently introduced crop plant, is the most susceptible plant species in the greenhouse and field, and the only plant species highly susceptible to infection by isolates from all three *E. tracheiphila* lineages. The high susceptibility of cucumber to isolates from all three clusters in both the field and greenhouse suggests that cucumbers could be functioning ecologically as a highly susceptible reservoir host. This presents the possibility that *E. tracheiphila* (which was already present in the Midwestern US by 1900 (66, 67)) could not have emerged or persisted as an annual epidemic without the human-mediated introduction of cultivated *Cucumis* spp. into temperate North America in the early 1500s (33).

*E. tracheiphila* has among the most dramatic structural genomic changes – including gene decay through pseudogenization, mobile element invasion and proliferation, and horizontal gene acquisitions – of any bacterial pathogen (20). These structural changes are consistent with a recent evolutionary transition from a progenitor with multiple environmental reservoirs and diverse metabolic capabilities to a pathogen with a narrow, host-specialized ecological niche. However, the species identity, geographic origin, and host relationships of the direct *E. tracheiphila* progenitor are all unknown, limiting our ability to investigate the evolutionary transition from the *E. tracheiphila* direct progenitor – presumably a plant commensal or weak pathogen – to a virulent pathogen (68-70). The genomic evidence of *E. tracheiphila*’s recent transition to a virulent, host restricted pathogen (20) highlights the continuing risk of non-pathogenic environmental microbes acquiring virulence genes via continual and naturally occurring mobile DNA invasion (71). Virulent pathogens are unlikely to persist in ecologically intact habitats with higher plant species diversity and higher diversity of pathogen resistance (R) genes within species (72-74). When pathogens evolve or acquire novel virulence genes, this acts as a selective pressure on host plant populations, and causes a rise in frequency of plant resistance genes. However, repeatedly planting the same crop plant varieties in agricultural populations interferes with this co-evolutionary dynamic by preventing a rise in frequency of effective host plant resistance. Identifying cultivars or wild crop relatives with resistance genes, and crossing them into cultivated crop populations, is one method favored by plant breeders. However, the probability of success from this approach for controlling *E. tracheiphila* is likely to be low. Cucumber is the best characterized of all cucurbit crops, and this species was found to contain among the lowest genetic heterogeneity of any vegetable crop, with an estimated effective population size of only 500 individuals at the time of domestication (75, 76). The *E. tracheiphila*-cucurbit association is evolutionarily novel (20), suggesting genetic resistance to *E. tracheiphila* may not exist in any undomesticated cucurbit populations. *Even if* the genetic basis of host resistance is identified in wild relatives or rare cultivars of cucumber, squash, or melon and successfully introduced into agricultural populations, *E. tracheiphila* is amenable to invasion by mobile DNA, including acquisition of virulence effector genes (20). This could function to quickly overcome potential host plant genetic resistance, especially if the same resistance gene(s) are broadly deployed in large, homogeneous crop plant populations (77, 78). This potential to rapidly generate novel variants from a recombining source population(s), together with ability to horizontally acquire virulence effectors will be important to consider when attempting to design durable resistance strategies for agricultural systems (20, 79).

Many – perhaps most – of the economically damaging plant pathogens and insect pests have emerged after the Neolithic Revolution (11, 16, 80-87). Yet, little effort has been put towards using ecological principles to plan genetic, physiological and/or structural complexity into agricultural systems to mitigate susceptibility to rapid spread of newly emerged insect pests or microbial pathogens (10). We hypothesize that the kind of local pathogen (or insect pest) emergence such as what has happened with *E. tracheiphila* is more common than currently understood. Further, we predict that these local emergence events can in some cases be followed by rapid dissemination through genetically homogeneous agricultural populations. Given the potential of such infections to threaten globalized crop populations, including staple crops that are vital for local and global food security, we urgently need to develop approaches for building sustainable agro-ecosystems that are rooted in ecological and evolutionary principles.

## Methods

### Study System

Wild species in the gourd family, Cucurbitaceae, occur in tropical and subtropical regions worldwide, and cultivars from this family are among the world’s most widely grown fruit and vegetable crops (34, 88). Like many Cucurbitaceae, *Cucurbita* spp. and *Cucumis* spp. produce a class of secondary metabolites called ‘cucurbitacins’ (89-91). Cucurbitacins are among the most bitter and toxic compounds ever characterized, and function as highly effective herbivory deterrents for almost all insect and mammalian herbivores, including humans (91-94). The exception are few genera of highly co-evolved Luperini leaf beetles (Coleoptera: Chrysomelidae), and for these beetles, cucurbitacins function as arrestants and feeding stimulants (92, 95, 96). *Acalymma* ssp. is a strictly New World genus of highly specialized leaf beetles that has co-evolved in Mesoamerica with *Cucurbita.* In natural settings, *Acalymma* spp. are obligately dependent on *Cucurbita* plants in all life stages (97-99). *E. tracheiphila* has no known environmental reservoirs, and only persists within infected *Cucurbita* spp. or *Cucumis* spp. plants, or the digestive tracts of the highly specialized beetle vectors. Beetle vectors are the only documented winter reservoirs of *E. tracheiphila* (45, 91, 100). The Eastern striped cucumber beetle (*Acalymma vittatum*) is the only species that has received substantial research attention because of its status as an important agricultural pest and plant pathogen vector in Eastern North America (99). *A. vittatum*, which is the predominant insect vector of *E. tracheiphila*, only occurs in Northeastern and Midwestern North America. It is likely that *A. vittatum* only recently emerged into this geographic area following the domestication and range expansion of *Cucurbita* for agriculture, as was recently shown for the obligate pollinator of *Cucurbita* in Eastern North America (65, 101). In the Old World, *Aulocophora* ssp. (Coleoptera: Chrysomelidae: Luperini) are obligate cucurbit specialists, although natural history information is almost completely absent for almost all species (102, 103).

### Confirmation of restricted *Erwinia tracheiphila* geographic range

Losses from *E. tracheiphila* are an annual epidemic in temperate Eastern North America (22, 25, 26, 29, 41, 89, 100, 104-108). No losses from *E. tracheiphila* have been reported anywhere else in the world. To evaluate whether the reported geographic restriction of *E. tracheiphila* to temperate Eastern North America is a reflection of its actual geographic range or an artifact of this pathogen not being recognized outside of this range, one of us (LRS) undertook extensive scouting expeditions of wild and cultivated *Cucurbita* spp., *Cucumis* spp., *Luffa* spp., and *Lagenaria* spp. populations in diverse areas of the world, including the entire Southern United States from California to South Carolina; on the west coast of Mexico from Jalisco to Oaxaca; in Europe, and in Southeast Asia. There is one report of *E. tracheiphila* in New Mexico (109), but this isolate was said to be from a cultivated watermelon (which is not susceptible) and this isolate is not archived nor do gene sequences from it exist, and we must therefore at this time consider it as a single erroneous report.

No *E. tracheiphila* symptoms were observed in undomesticated populations of *Cucurbita digitata* in California and Arizona, undomesticated or domesticated *Cucurbita* spp. or *Cucumis* spp. in California, Arizona, New Mexico, Texas, Louisiana, Mississippi, Alabama, Georgia, South Carolina, or Missouri. In Mexico, *E. tracheiphila* was not found in wild or cultivated cucurbits in the Mexican states of Jalisco, Guerrero, Michoacan, Oaxaca, Guanajuato, or Querétaro. Nor was *E. tracheiphila* observed in any cucurbits in commercial or academic farms in Thailand, the Philipines, or Vietnam. In Europe, *E. tracheiphila* was never observed in cucumber or squash plants in Spain or Germany. These observations are consistent with the lack of reports of *E. tracheiphila* outside of temperate Northeastern and Midwestern North America. *E. tracheiphila* has never been shown to survive outside of few agricultural species of cucurbit hosts and beetle vectors (45, 100). Therefore, the isolates collected in this study (Figure 2a, Figure 2b, Supplemental Table 1) are hypothesized to cover the entire plant host and geographic range where *Erwinia tracheiphila* exists.

### Collecting single isolates of *Erwinia tracheiphila*

Single *E. tracheiphila* isolates were obtained from symptomatic squash (*Cucurbita* spp.) muskmelon (*Cucumis melo*), and cucumber (*Cucumis sativus*) plants in agro-ecosystems from across the entire geographic range where economic losses from *E. tracheiphila* are reported (Supplemental Table 1, Figure 2b). In the field, infected plants were visually identified by characteristic wilting symptoms (Figure 1a). All wilting, symptomatic plants in a given field were gathered to avoid collection bias. Symptomatic vines from infected plants were removed with a sterile knife, immediately placed in separate 1 gallon plastic bags, and stored at 4°C for a maximum of 3 days prior to performing bacterial isolations. The reference BuffGH strain (formerly PSU-1) was isolated in 2007 from an undomesticated wild gourd *C. pepo* ssp. *texana* plant growing at the Rock Springs Experimental Station in Rock Springs, Pennsylvania. These *C. pepo* ssp *texana* seeds were originally collected from wild populations in New Mexico and Texas, and are now greenhouse cultivated then field transplanted for academic research at Pennsylvania State University in University Park, PA (reviewed in (89)). Isolates collected in 2007-2009 were acquired from (51), collected according to the protocol described therein, and are stored at Iowa State University in Ames, IA. *E. tracheiphila* isolates from 2015 were collected by first washing external dirt and debris from symptomatic vines with tap water, and then surface sterilizing the cleaned vines with 70% ethanol. Sterilized vines were cut into 3-4 inch sections between nodes with sterile razor blades, and ½ inch of the vine sections were soaked in 3 ml of autoclaved Milli-Q water in 15ml Falcon tubes until pure *E. tracheiphila* could be seen on the cut surface (Figure 1b). Sterile loops were then used to transfer *E. tracheiphila* ooze (Figure 1b) to King’s B agar plates (1L: 20g protease peptone #3, 10ml glycerol, 1.5g MgSO_4_•7H_2_O; 1.5g KH_2_PO_4_; 15g bactoagar). Single isolates were restreaked, and then single colonies from the restreaked plates were grown in shaken liquid KB broth at 25°C for 48 hours and cryogenically preserved with 15% glycerol.

### DNA extraction, library preparation and whole genome sequencing

Single colonies from cryogenically preserved glycerol stock were grown on KB agar plates, and single colonies were grown in liquid KB for 36-48 hours, or until OD_600_ = 1. DNA from liquid cultures was extracted with Promega DNA wizard (Promega, Madison, WI) following manufacturer’s instructions.

Libraries of the genomic DNA for isolates listed in Supplemental Table 1 were generated using a Nextera DNA Sample Preparation Kit (Illumina, San Diego, CA). The libraries were amplified for 8 cycles using the KAPA HiFi Library Amplification Kit (KAPA Biosystems, Wilmington, MA), and the size selection was performed using AMPure XP beads (Agencourt Bioscience Corp, Beverly, MA). Library concentrations were measured using a QuBit DNA Quantification Kit (Life Technologies, Carlsbad, CA) and the fragment size range detection (100-400bp) was performed using the TapeStation 2200 (Agilent Technologies, Santa Clara, CA). Libraries were pooled using Nextera Index kits and 150-bp paired-end reads were generated with an Illumina HiSeq 2500 Sequencing System. Assembly metrics of all strains sequenced for this study were determined with QUAST, with standard settings that only retain contigs larger than 500bp (110).

### Transformation of *Erwinia tracheiphila* with an mCherry expressing plasmid

*E. tracheiphila* strain BuffGH was used for visualization of *E. tracheiphila* in the xylem of infected squash seedlings. Plasmid pMP7605 carrying a constitutively expressed mCherry gene was electroporated into competent *E. tracheiphila* cells. For this, we followed protocols described previously (111). Briefly, competent *E. tracheiphila* cells were prepared by growing *E. tracheiphila* in 200 ml KB to OD_600_ = 0.02. Subsequently, cells were washed using decreasing volumes, once with chilled sterile milliQ water, twice with 10% glycerol, and finally resuspended in 2 ml of 10% glycerol. For electroporation, a 40 μl aliquot of competent cells was mixed with 4 μl of plasmid DNA, placed in a 0.2 cm cuvette and electroporated at 2.5 kV for 5.2-5.8 ms. Electroporated cells were immediately transferred to 3 ml KB liquid and incubated at room temperature without shaking for 1 hour. A cell pellet was obtained and resuspended in 100 μl of media, and then plated in KB agar with ampicillin (100 μg/ml). Colonies of fluorescent *E. tracheiphila* were obtained after 5 days at room temperature.

### Genome Assembly and Annotation

Adaptor trimming and quality filtering of raw Illumina reads was performed using the FastX toolkit 0.0.13.2 (Pearson, et al. 1997), SeqTK 1.0 (https://github.com/lh3/seqtk/), and FastQC 0.10.1 (http://www.bioinformatics.bbsrc.ac.uk/projects/fastqc/). Both mapping and *de novo* assemblies were then generated for each sequenced isolate. For the *de novo* assemblies, SPAdes 3.1.1 was used with default parameters to assemble the quality filtered, adapter trimmed, paired end reads using k-mer sizes of 21, 33, 55 and the –careful parameter (112). For *ab initio* annotations of the assembled *de novo* whole genome sequences, PROKKA version 1.11 was used with default parameters (113). For the mapping-based assemblies, Mira 4.1 (114) was used to map quality filtered, adapter trimmed, paired end reads from each isolate to the BuffGH PacBio reference strain (30). The functional annotations of all coding sequences (including pseudogenes) were transferred to each genome from the manually-curated annotation of the reference BuffGH genome (20), using the RATT function in PAGIT 1.0 (115). We assumed that all pseudogenes are the same in all isolates, which will only be confirmable with long read PacBio sequencing of these isolates followed by manual annotations.

### Phylogenetic relationships of *Erwinia tracheiphila* isolates

Orthologous gene families present in all *E. tracheiphila* isolates were identified from the *de novo* assemblies with OrthoMCL (116) through an all-versus-all BLASTP 2.2.28 + search with an *E*-value cutoff of 10^−5^. The orthologous genes were aligned using MAFFT 6.853 (117). The gene alignments were trimmed with trimAl version 1.2 using the “automated1” option (118). The individual gene alignments were concatenated into the core genome alignment using the publically available script (https://github.com/tatumdmortimer/core-genome-alignment, last accessed September 8, 2014). The 237,634 concatenated core genome alignment used to reconstruct the network analysis in Figure 2a is included as Supplemental File 1. The evolutionary relationships among the isolates were reconstructed and visualized in SplitsTree v 4.13.1 (47) using the core genome alignment as input.

### Determination of within-cluster diversity

The genes from the manual annotations transferred to the mapped assemblies were used in an all-versus-all BLASTP 2.2.28 + (119) search with an *E*-value cutoff of 10^−5^. OrthoMCL (116) was run separately for all the isolates within each lineage to identify the core orthologous gene families within each of *Et-melo, Et-C1*, and *Et-C2*. For population genetics analyses, the ‘core’ genes shared by all isolates within each of the three lineages were designated as either ‘*Intact’*, meaning they are putatively functional based on the manually curated annotations in (20), or ‘*Pseudogenized/Repetitive’*, meaning they are either predicted to be pseudogenes or were predicted to be mobile DNA (genes from bacteriophage, insertion sequences, plasmids, or transposases). The ‘*Pseudogenized/Repetitive’* genes from bacteriophage, insertion sequences, plasmids, or transposases were determined by domain assignments with PfamScan 1.5 (120), ISFinder (January 2015 update) (121), and PHAST (122) as described in (20). For *Et-C1* and *Et-melo* clusters sampled at multiple time points, two groups were created: isolates collected from 2008-2010 or collected in 2015. Genetic diversity was quantified for each cluster using Watterson’s estimator, *θ_W_* per site (123), where *θ_W_* estimates 2*N_e_μ*, where *N_e_* is the effective population size and *μ* is the mutation rate.

For recombination estimates, quality filtered reads were mapped to the reference BuffGH sequence (30) with Burrows-Wheeler Alignment (BWA) tool 0.7.4 (124), a pileup was created with SAMTools 0.1.18 (125), and variants were called with VCFtools 0.1.9 if the phred quality score of the variant site was greater than or equal to 60 (126). Single nucleotide polymorphisms (SNPs) were not called if (i) within 9 base pairs (three codons) of each other and (ii) with less than 10X coverage or (iii) with more than 150X coverage, since short Illumina reads cannot be accurately placed over repetitive regions. Recombination rates within each pathovar were estimated by using the ‘VCF_to_FASTA.sh’ (Supplemental File 2) script to create whole genome alignments compatible with gubbins 2.1.0 (127), which was run for standard 10 iterations.

### Pangenome Identification

The Micropan package (128) in R 3.2 (129) was used to identify the core and pangenome of *de novo E. tracheiphila* isolate assemblies. *De novo* assemblies (see ‘Genome Assembly and Annotation’ section above) were used to ensure that the entire repertoire of genes present per isolate were included, and the pangenome estimates would not be biased with mapping assemblies based on what was present in the reference genome. The groups.txt output file from the OrthoMCL clustering of protein sequences of the *de novo* assemblies (see ‘Phylogenetic relationships of *Erwinia tracheiphila* isolates’ section above) and custom R scripts (129) were used to identify genes that were ‘rare’ (present in less than 5% of isolates) or ‘core’ (present in more than 95% of the sequenced isolates).

### Functional comparison of core and rare genes

The *ab initio* predicted genes from each *E. tracheiphila* sequenced isolate were searched against the Clusters of Orthologous Groups (COG) database (2014 update; Galperin et al. 2014) using BLASTP 2.2.28+ (Altschul et al. 1990). Only the top-scoring match (per gene) with *E*-value <10^−5^ was kept. Each gene was assigned a COG category of the first functional category of the top-scoring match. Genes without significant matches to any sequence in the COG database were not assigned a functional category. A one-way Fisher’s exact test with corrections for multiple comparisons was used to identify the COG categories enriched in each cluster and graphed with ggplot2 in R (129).

### Identification of T3SS virulence genes and reconstruction of effector gene phylogenetic trees

The *ab initio* coding sequences predicted by PROKKA from each *E. tracheiphila* isolate were compared against a manually curated version of the *Pseudomonas* Hop protein effector database (http://www.pseudomonas-syringae.org/T3SS-Hops.xls; accessed August 28, 2015, with additional non-*Pseudomonas hrp*T3SS effectors manually curated; Supplemental File 3) using BLASTP with an e-value cutoff of 10^−5^. Presence and absence of effector genes was visualized with gplots (130) in R 3.2 (129).

To reconstruct the phylogeny of the cluster specific effector genes identified in *E. tracheiphila*, the amino acid sequence of each gene was used as a BLASTP query against the *nr* database (119). An e-value cutoff of 10^−5^ was used to acquire a phylogenetically representative sample of homologs. The sequences were aligned with MAFFT v. 6.853 (117) and trimmed with trimAl 1.2 (118). The maximum-likelihood phylogeny of each aligned gene was reconstructed using RAxML 8.2.4 (131) as implemented on the CIPRES server (132), under the GTR + CAT model and with 100 bootstrap replicates. Bootstrapped pseudoreplicates were summarized with sumTrees.py 4.0.0 (133), and the bootstrap consensus tree was visualized with FigTree 1.4.2 (134).

### Cross inoculation experiments

Seven *E. tracheiphila* isolates from the three different phylogenetic clusters were randomly chosen and used for testing virulence (i.e., the degree of harm) of isolates from each cluster against susceptible host plant species. From *Et-melo*, the experimental isolates were HPsat9.4, RHmelo4.4, RHsat4.1, from *Et-C1* the isolates were BHKY, HPcuc8.2, and PAsat11typeII, and from *Et-C2* the isolate was UVM3.1. Single colonies of each isolate were grown in liquid KB for 24 hours until mid-exponential phase, and then all strains were diluted to OD_600_ = 0.3. For the inoculations, 25μl of culture from each isolate was then applied to a small break in a single leaf petiole of two-week old seedlings at the two-leaf stage. Plants were observed several times weekly for the initial appearance of wilt symptoms in the inoculated leaf, spread of symptoms to a second leaf, and plant death within a 25 day experimental period, following (22, 25, 28, 43, 44, 106, 135). Plant death was scored when all leaves were determined to be too desiccated to support beetle vector herbivory, which is necessary for acquisition of *E. tracheiphila* by beetle vectors and subsequent transmission to healthy hosts. At this stage of infection, the leaves are also too desiccated for photosynthesis. For the statistical analysis, *Et-C1* and *Et-C2* inoculation data were not statistically different, and because they were not different and they share the same host plant range, these were combined. A one-way ANOVA with either ‘Days until first wilt symptoms’ or ‘Days until Death’ as the response and ‘Host plant species’ was conducted for both *Et-melo* data and *Et-C1* + *Et-C2* data with model statement aov(lm(Growth~Cluster) as implemented in R 3.2 (129).

### Effects of temperature on *in vitro* growth rate

Twelve *E. tracheiphila* isolates from the three different phylogenetic clusters were randomly chosen for testing the effect of temperature on *in vitro* bacterial growth.

The isolates from *Et-C1* are HPcuc8.2, PAsat3.1, PAsat2.3, and BHKY, from *Et-C2* are ConPepo4M2, ConPepo4M1, UVM3.1 and from *Et-melo* are RHmelo2.1, RHmelo4.4, RHsat4.1, PAsat11typeIII, HPsat9.4. The starting cultures were prepared by inoculating a single colony of each *E. tracheiphila* isolate into 4 ml of KB media, which was grown at room temperature with shaking for 48 h until stationary phase. One ml of each culture was then pelleted, washed with 1 ml of fresh KB media, and resuspended in the same volume. The washed cell suspensions were then diluted in fresh KB media to OD_600_=0.04, and then four 300 μl replicates of each diluted cell suspension were placed in a single well of an optically clear 96 well plate. The 96 well plate was placed in a plate reader (Spectra Max V2) and absorbance (OD_600_) was measured every 5h over a total 40h experimental period at 28°C, 30°C, 33°C, and 37°C, with non-inoculated KB media used as a negative control. A two-way ANOVA to test the effects of ‘Temperature’, ‘Cluster’, and their interaction term on the OD_600_ concentration at different temperatures the model statement OD_600_ = Temperature + Cluster + Temperature*Cluster as implemented in R (129).

### Effects of temperate and subtropical temperatures on *in planta* virulence

Single colonies of two *E. tracheiphila* isolates – one derived from a field-infected cucumber (HPcuc8.2, cluster *Et-C1*), and one from a field-infected squash (BHKY, cluster *Et-C1*) – were each grown in liquid KB to an overnight exponential phase concentration of OD_600_=0.3. The squash origin isolate was inoculated into two-week old squash seedlings at the two leaf stage, and the cucumber origin isolate was inoculated into two week old cucumber plants at the two leaf stage. All plants were inoculated with 25µl of bacterial inoculum placed on a single petiole wound. The seed varieties used were *Cucurbita pepo* ‘Dixie’ squash and *Cucumis sativus* ‘Marketmore’ cucumber from Johnny’s seeds (http://www.johnnyseeds.com/).

Average July temperatures for Texas and Massachusetts were determined by a Google search to be 33°C day and 23°C nights for Texas, and 27°C day and 18°C night for Massachusetts. All plants were kept in programmable Conviron growth chambers with 16hr light/8hr dark, 60% relative humidity. Plants were observed several times weekly for the initial appearance of wilt symptoms in the inoculated leaf, spread of symptoms to a second leaf, and plant death within a 25 day experimental period, following (22, 25, 28, 43, 44, 106, 135). The sample sizes used in this experiment are n=44 Texas ‘Dixie’ squash, n=44 Massachusetts ‘Dixie’ squash, n=34 Massachusetts ‘Marketmore’ cucumbers, and n=36 Texas ‘Marketmore’ cucumbers. A one-way ANOVA to test the effects of ‘HostSpecies’ (either cucumber or squash) at both ‘Texas’ and ‘Massachussetts’ temperatures with the model statements *Death = State + HostSpecies*; and *Wilt = State + HostSpecies* as implemented in R (129).

### Data Deposition

Raw reads from the sequenced isolates (Supplemental Table 1) are available at the NCBI BioProject PRJNA272881. The sequence filtering and analysis pipeline, micropan parameters for pangenome analysis, modified Hop *hrp*T3SS database, and ‘VCF_to_FASTA.sh’ script used to create FASTA alignments of variant calls for recombination analysis in gubbins, will be available via FigShare at the time of manuscript acceptance.

## Acknowledgements

This study was made possible by an NSF postdoctoral fellowship DBI-1202736 to LRS; NIH Grant GM58213 to R.K.; OZ was in part supported by a Simons Investigator Award from the Simons Foundation; computational advice and support from, Bob Freedman and Aaron Kitzmiller at Harvard FAS; Harvard Odyssey Computational Resources; Taj Azarian for helpful advice and discussion; Scott Chimileski for Figure 1b image; Rob Dunn for constructive comments on the manuscript; Weld Research Building at the Arnold Arboretum for staff support, confocal microscope access, and growth chamber facilities; Miguel Coehlo for Nextera library preparation help; Eric Alm for providing the original suggestion to sequence dozens of isolates; infected plant samples provided from Christian Herter Community Garden; Verrill Farms, Dana Roberts, Kristy’s Barn, the University of Vermont Horticultural Farm, Green Valley Farm, and all other farms and individuals that contributed isolates; Nora Mishanec and Laura Jenny for laboratory and greenhouse assistance. The US Department of Agriculture, Agricultural Research Service, is an equal opportunity/affirmative action employer and all agency services are available without discrimination. Mention of commercial products and organizations in this manuscript is solely to provide specific information. It does not constitute endorsement by USDA-ARS over other products and organizations not mentioned.

## Author Contributions

LRS conceived of the study; RK supervised the development of the project, LRS, JNP, BJA, EDS, and KH performed computational analysis; LRS and JR did laboratory and greenhouse experiments; all authors contributed to the interpretation of data; LRS drafted the original article and JNP, BJA, EDS, OZ, NP, JR, VKC, KH, and RK contributed revisions.

---

**Supplemental Figure 1. The phylogenetic network of 88 *Erwinia tracheiphila* isolates.** Isolates are identified by number, and all associated metadata are listed in Supplemental Table 1.

**Supplemental Figure 2. The distribution of virulence effector genes in 88 *Erwinia tracheiphila* isolates.** Each individual sequenced isolate is represented by a row, and the rows are grouped by phylogenetic cluster (y axis). The 23 effector genes found in the *Erwinia tracheiphila* pangenome are arranged on the x axis. Each cell in the matrix is color-coded by the presence (blue) or absence (dark grey) of a *hrp*T3SS effector gene in a genome of an individual isolate. Isolate ID metadata are listed in Supplemental Table 1. Isolates are identified by number (far right axis) that corresponds with ID in Supplemental Table 1.

**Supplemental Figure 3.** Unrooted maximum likelihood phylogenetic trees of the five cluster-specific effector genes (HopAN1, NleD, AvrRpm1, Eop3, HopAF1). The tree topologies of these genes are suggestive of horizontal acquisition by *E. tracheiphila*, as these effector genes are sparse or absent in other plant-associated *Erwinia* spp. and in the sister genus *Pantoea* spp. Bootstrap support values greater than 50% are shown at the nodes. Scale bar, number of substitutions per site.

